# Effect of gastrointestinal alterations mimicking elderly conditions on in vitro digestion of meat and soy proteins

**DOI:** 10.1101/2021.11.29.470321

**Authors:** Chong Wang, Fan Zhao, Yun Bai, Chunbao Li, Xinglian Xu, Karsten Kristiansen, Guanghong Zhou

**Affiliations:** College of Food Science and Technology, Nanjing Agricultural University; Key Laboratory of Meat Products Processing, Ministry of Agriculture; Jiangsu Collaborative Innovation Center of Meat Production and Processing, Quality and Safety Control, Nanjing, 210095, P.R. China; Laboratory of Genomics and Molecular Biomedicine, Department of Biology, University of Copenhagen, Copenhagen 2100, Denmark; Institute of Metagenomics, BGI-Shenzhen, Shenzhen 518083, P. R. China

**Keywords:** Elderly, Protein sources, In vitro digestion, Digestibility, Proteomics

## Abstract

We evaluated the digestion of meat (chicken, beef, and pork) and soy proteins under in vitro conditions mimicking gastrointestinal (GI) conditions of adults (control, C) and elderly with achlorhydria (EA). The changes in degree of hydrolysis (DH), SDS-PAGE profiles, peptide concentration, and proteomics profiles during the digestion process were investigated. Digestion under the EA condition markedly decreased the DH of all protein sources, especially for soy protein. SDS-PAGE profiling and proteomics showed that myofibrillar/sarcoplasmic protein from meat and glycinin/beta-conglycinin from soy were the proteins most affected by the different digestive conditions. Our results indicated that the difference in the digestibility of meat protein between EA and control conditions gradually narrowed from the gastric to the intestinal phase, while a pronounced difference between control and EA conditions was maintained also in the intestinal phase. This work provides new insights of value for future dietary recommendations for elderly individuals.

## 1. Introduction

Life expectancy has increased worldwide and the population aged more than 60 years old will reach two billion by 2050 (Nations, 2019). According to the recent Chinese data of the seventh national population census, the number of individuals aged 60 years or beyond surpasses infants and youth (aged below 14 years of age) (Cheng & Duan, 2021). Consequently, this will be accompanied by global major challenges in relation to elderly wellness, including lifestyle and nutritional issues. The physiological functions declining with aging include gastrointestinal (GI tract) functions (Shani-Levi et al., 2017) with achlorhydria - the absence of gastric acid (hydrochloric acid) secretion in the stomach being one of the most common age-related gut disorders. The prevalence of achlorhydria increases from 2.5% in persons in the 30s to 12% in persons in the 80s (Villanacci et al., 2017). A study involving 3484 subjects revealed that 27% suffered from a varying degree of achlorhydria, and the greatest incidence (39.8%) occurred in females 80 to 89 years of age (Sharp & Fister, 1967). Gastric acid plays a pivotal role in food digestion by activating pepsin. Simultaneously, the acidic environment also inhibits or eliminates many microorganisms. Accordingly, achlorhydria is likely to influence digestion and absorption of food constituents in a number of elderly individuals. An in vitro study on gastric digestion of major milk proteins indeed revealed that hydrolysis of caseins proceeded slower in conditions mimicking digestion in elderly compared to those of younger individuals (Aalaei, Khakimov, De Gobba, & Ahrné, 2021).

Food composition, protein sources and amino acid composition, physicochemical characteristics including texture, structure and solubility affect the bioavailability of proteins. Wen et al. (2015) reported that pepsin-mediated in vitro digestion of pork and beef produced similar patterns of peptides that differed from peptides produced by digestion of chicken and fish meat, and clear differences in the rate of digestion of casein and whey have been well documented (Dangin et al., 2001). However, these digestion studies were not performed using conditions mimicking the conditions in the stomach of elderly, and thus, studies exploiting stomach conditions resembling the conditions in elderly are warranted, also in order to provide dietary recommendations for elderly individuals.

Due to high cost and technical and ethical restrictions of animal studies, in vitro digestion has become an alternative method to evaluate digestibility of proteins enabling easy reproducible sampling (Hur, Lim, Decker, & McClements, 2011). Thus, an international consensus for a static protocol, INFOGEST, reflecting the physiological conditions in humans was adopted in 2014 (Minekus et al., 2014). Recently, in vitro digestion studies using conditions resembling those of the GI tract in elderly have been reported. Hernández-Olivas, Muñoz-Pina, Andrés, & Heredia (2020) investigated the effect of altered GI conditions on the proteolysis of different kinds of fish (salmon, sardine, sea bass and hake). They found that proteolysis was significantly affected using conditions mimicking the alterations observed in the elderly GI (33-42% reduction). Subsequently, this research group also assessed the impact of cooking of eggs (hard-boiled, poached, and omelet) on protein digestion, and based on the results they recommended intake of hard-boiled and poached eggs to elderly individuals (Hernández-Olivas, Muñoz-Pina, Andrés, & Heredia, 2021). Even though these studies have indicated impaired digestion in elderly, details on how different protein sources are digested in elderly versus young individuals are still scarce, and more detailed studies are clearly warranted.

The present study applied in vitro digestion models mimicking young adult and elderly conditions to investigate the digestion profiles of meat (chicken, beef and pork) protein and soy protein, focusing on elucidating protein digestion changes under altered digestion conditions.

## 2. Materials and methods

### 2.1 Materials

Pepsin from porcine gastric mucosa (≥2500 U/mg), pancreatin from porcine pancreas (8 × USP), bovine bile powder, analytical grade salts (CaCl_2_, KCl, KH_2_PO_4_, NaHCO_3_, NaCl, MgCl_2_(H_2_O)_6_, (NH_4_)_2_CO_3_, CaCl_2_(H_2_O)_2_), NaOH, HCl, methanol (HPLC grade, ≥99.9%), dichloromethane (HPLC grade, ≥99.9%), acetone (HPLC grade), trifluoroacetic acid, L-leucine and fluorescamine were purchased from Sigma-Aldrich (Deisenhofen, Germany). Chicken pectoralis major muscle, beef longissimus dorsi muscle, pork longissimus dorsi muscle, and soy protein (isoflavones were removed by extraction with 80% methanol) were purchased from Linyi Shansong Biological Products Co., Ltd. (Linyi, China).

### 2.2 Sample preparation

After removal of the visible fat and connective tissues, the muscles were cut into pieces, placed in plastic bag and cooked in a 90 °C water bath until the internal temperature reached 70 °C. After cooking, meat samples were cooled at room temperature and ground into powder. The intramuscular fat was extracted and removed by using a mixture of methylene chloride/methanol (2:1, v/v), the extraction process was repeated once. After fat extraction, the residual solvent was removed by evaporation and the resulting protein powder was passed through a 100 Mesh sieve. The final protein powders were stored at −20 °C.

### 2.3 In vitro digestion

A static in vitro digestion system was applied. The control model corresponding to standard GI conditions of healthy young adults was established according to the standardized INFOGEST protocol (Minekus et al., 2014). The elderly model corresponding to conditions of the elderly population with achlorhydria was established according to Shani-Levi et al. (2017) and Russell et al. (1993). The specific conditions of each model are shown in Suppl. Table 1. Briefly, protein powder (1.0 g) was mixed with 14.4 mL simulated gastric fluid (SGF), and then this mixture was homogenized. After that, 6 mol/L HCl were added to adjust the pH to the predetermined values, 3.0 for control and 6.0 for elderly conditions, respectively, and 1.6 mL pepsin solution were added to start the gastric digestion. The mixture was incubated at 37 °C for 2 h. After 1, 10, 20, 30, 60 and 120 min, 0.5 mL of digest was collected and mixed with 0.5 mL of simulated intestinal fluid (SIF) to inactivate pepsin. For intestinal digestion, 10 mL of gastric digest were withdrawn and mixed with 6 mL SIF. After pH adjustment to 7.0, 1.25 mL bile and 2 mL pancreatin solution were added to start the intestinal digestion. The mixture was incubated at 37 °C for 2 h or 4 h depending on the digestive model. After 1, 10, 20, 30, 60, 120 and 240 min, 0.5 mL of digestive solution was withdrawn and pefabloc was added to a final concentration of 4 mM to stop intestinal digestion. All aliquots were stored at −80 °C prior to further analysis.

### 2.4 Degree of Hydrolysis

A fluorecamine assay was carried out to evaluate the degree of hydrolysis (DH) of digested protein as described by (Larsen, Rasmussen, Bjerring, & Nielsen, 2004). Briefly, 75 μL of each digesta were mixed with 24% trichloroacetic acid (v/v=1:1) and precipitated on ice for 30 min. The solution was centrifuged at 4 °C for 30 min at 13000 rpm. Afterward, 30 μL of standard (L-leucine) or digesta were mixed with 900 μL sodium tetraborate (0.1 M, pH 8.0) and 300 μL fluorescamine acetone solution (0.2 mg/mL). Fluorescence was determined at excitation and emission wavelengths of 390 and 480 nm, respectively. The DH was calculated as followed:

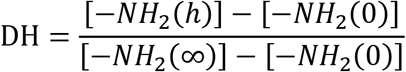

[-NH_2_ (h)] and [-NH_2_ (0)] denotes the concentration of primary amines in the hydrolyzed (h) and unhydrolyzed (0) samples, respectively. [-NH_2_ (∞)], indicating the theoretical maximal primary amine concentration measured by fluorescence of a raw sample which was completely hydrolyzed by HCl at 100 °C for 48 h.

### 2.5 SDS-PAGEs

Proteins in the digesta were separated and quantified by SDS-PAGE with 4-20% precast gel (Genscript, USA). Each sample was diluted and mixed with 5× loading buffer to reach the concentration of 4 mg/mL. All samples were heated at 95 °C for 5 min and then loaded onto the gel. The gels were run at 180 V for approximately 40 min until the band of bromophenol blue dye just disappeared. After electrophoresis, the gels were stained by Coomassie Brilliant Blue R250 for 0.5 h and destained until the bands were clear. Gel images were obtained by using an image scanner (GE Healthcare, U.K.)

### 2.6 LC-MS/MS analysis of digested proteins

A label-free proteomics protocol was applied to determine protein profiles. All samples were centrifuged at 12000 rpm for 20 min at 4 °C, and the supernatant was filtered using a 10 kDa molecular weight cut-off ultrafiltration tube (Millipore, USA). After that, the peptide mixtures were desalted using C18 cartridges (Waters, USA), concentrated by vacuum centrifugation and reconstituted with 0.1% TFA. The peptide concentration was assessed using Nanodrop at 280 nm.

A Nano-LC 1000 tandem with Q Exactive mass spectrometer (Thermo Fisher Scientific, USA) was used to separate and analyze the peptide profiles. 2 μL of each sample were loaded onto a RP-C18 column (15 mm × 75 μm × 3 μm, Thermo Fisher Scientific). 2% acetonitrile with 0.1 % formic acid (A) and 80% acetonitrile with 0.1 % formic acid (B) were used for mobile phases, and the flow rate was 250 nL/min. The gradient program was as follows: 0-50 min, linear gradient from 4 to 50% eluent B; 50-54 min, linear gradient from 50-100% eluent B; 54-60 min, 100% eluent B. Electrospray ionization (ESI) was applied in the positive mode with the following parameters: MS data was acquired using a data-dependent top 10 method dynamically exclusion to screen the most abundant precursor ions from survey scan (300-1800 m/z) for HCD fragmentation. Dynamic exclusion duration was 25 s. Survey scans were acquired at a resolution of 70,000 at m/z 200 and resolution for HCD spectra was set to 17,500 at m/z 200. Normalized collision energy was 30 eV.

The MS data were analyzed using the MaxQuant software (version 1.3.0.5) and searched against the corresponding UniProt *Gallus gallus*, *Bos Taurus, Sus scrofa, Glycine max* database. The precursor mass and MS/MS tolerance of peptide were set to 6 and 20 ppm, respectively. No enzyme was selected to conduct enzymatic cleavage. The cut-off global false discovery rate (FDR) for peptide and protein identification was set to 1%. Protein abundance was calculated on the basis of the normalized spectral protein intensity (LFQ intensity). The protein intensity was quantified based on the razor and unique peptides intensity.

### 2.7 Statistics analysis

All results were calculated as the means and standard deviations. The results were analyzed by ANOVA (p<0.05). Comparison of the mean values was performed using Duncan’s test. Statistical analyses were performed with SPSS for Windows version 20 (SPSS Inc., Chicago, IL).

## 3. Results and discussion

### 3.1 In vitro digestibility

The digestibility of different proteins under control and EA conditions was compared by measuring the liberation of free amino acid using the fluorescamine method. The effect of alterative digestive conditions on DH is shown in Figure 1. The DH of all protein groups increased with digestion time, but exhibited differences in relation to both digestive phases and protein sources. In the gastric phase, compared with the control condition, the elderly achlorhydria (EA) digestion condition overall led to a significant decrease in DH values regardless of protein sources. This result is consistent with Hernández-Olivas et al. (2020), who found that the proteolysis of fish in condition mimicking elderly GI conditions was lower (33-50% reduction) than using condition mimicking young adults. Of note, the digestibility of soy protein in the control condition was significantly higher than that of meat proteins, while digestibility of soy protein was the lowest when digested using the EA condition.

**Figure 1.**
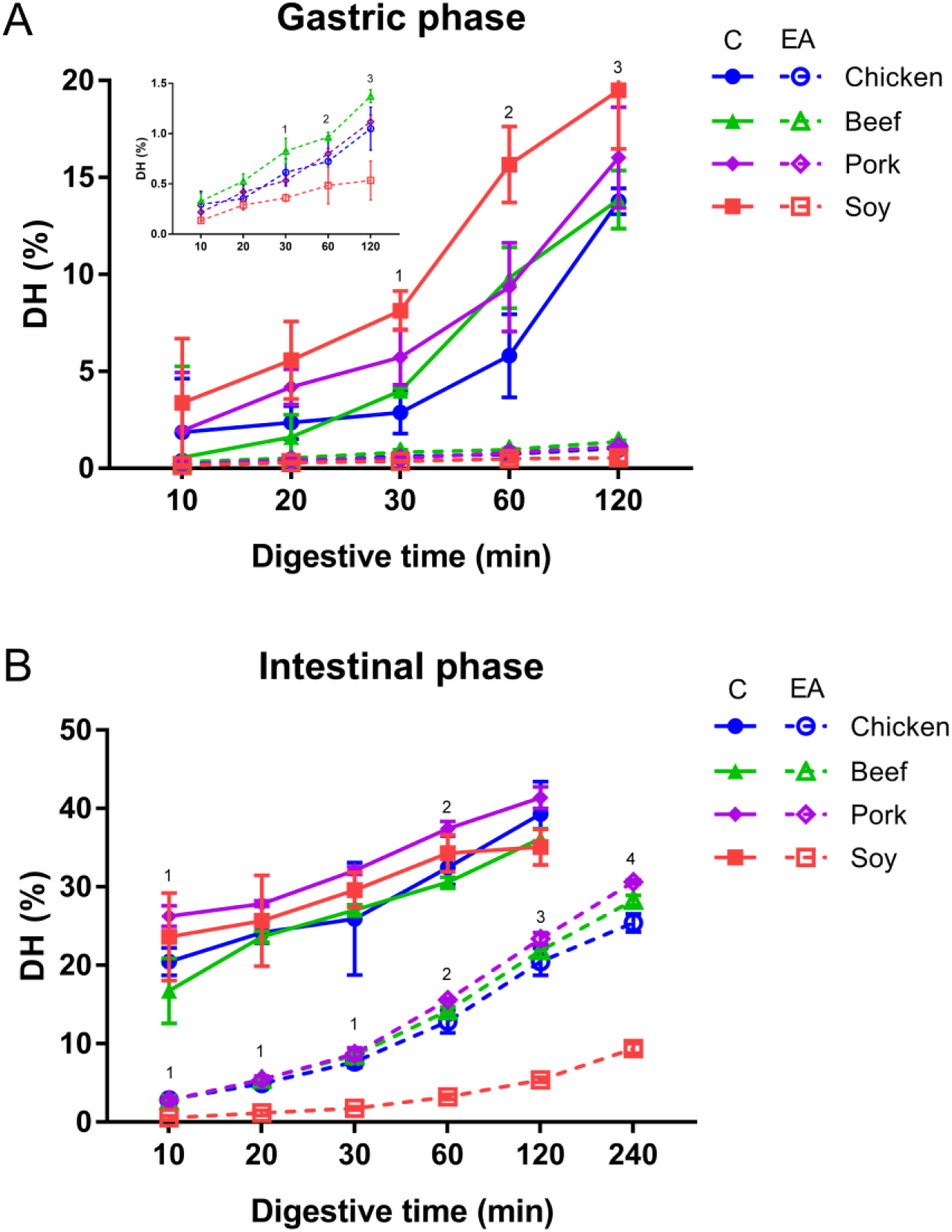
Degree of hydrolysis of different proteins in the gastric phase (A) and the intestinal phase (B). Meat and soy proteins were subjected to in vitro digestion using conditions that mimicked the gastro-intestinal conditions in young healthy individuals (control, C) and elderly individuals with decreased secretion of HCl in the stomach, achlorhydria (EA). The degree of hydrolysis (DH) was followed using a fluorecamine-based assay. Error bars represent the standard deviation obtained from three replicated experiments. Statistical analysis (*p* <0.05): Control gastric phase (1: chicken vs soy; 2: chicken, beef, pork vs soy; 3: chicken, beef vs soy). EA gastric phase (1: chicken, beef vs soy; beef vs pork; 2: beef, pork vs soy; 3: chicken, beef, pork vs soy; beef vs pork; chicken vs beef). Control intestinal phase (1: beef vs pork; beef vs soy; 2: beef vs pork). EA intestinal phase (1: chicken, beef, pork vs soy; 2: chicken, beef, pork vs soy; chicken vs pork; 3: chicken, beef, pork vs soy; chicken, beef vs pork; 4: chicken, beef, pork vs soy; chicken, beef vs pork; chicken vs beef).

According to Shani-Levi et al. (2017), the pepsin level in the stomach of the elderly is reduced by 25% compared to that of young adults. Hence, we reduced the pepsin concentration for the EA condition from 2000 to 1500 U/mL, which resulted in a notably lower digestibility of all proteins under the EA condition compared to the control condition. The activity of pepsin, the major protease in the stomach, is highly affected by the pH of the gastric juice (Roberts, 2006), and few studies have reported on the increased pH in the stomach of elderly with achlorhydria, and a slower return to the low gastric pH characterizing the fasting state after intake of a meal (Husebye, Skar, Høverstad, & Melby, 1992; Russell et al., 1993). In addition, low pH tends to result in a change in secondary or tertiary structure, resulting in the exposure of enzyme cleavage sites (Xiao et al., 2016). Carbonaro, Maselli, & Nucara (2012) found a strong negative correlation between the β-sheet structure and food digestibility, and an inverse linear correlation was observed between antiparallel β-sheet structures and protein digestibility. Wan & Guo (2019) demonstrated that soy protein adopts different conformation in relation to pH, including β-sheet aggregation and fibril formation. Thus, our results indicate that the digestive properties of soy protein compared to meat proteins are greatly affected by pH, implying an impaired ability to digest soy protein in elderly individuals.

As chyme gradually empties into the small intestine, pancreatic proteases are secreted with pancreatic bicarbonate. The optimum pH for key proteases of the small intestine is known to be 7.0-9.0 (Rick, 1974). Shani-Levi et al. (2017) reported that the intestinal pH of the elderly is similar to young adults. However, low levels of proteolytic enzymes (reduced to 50%) and bile are present in the small intestine of elderly compared to young adults. Accordingly, the digestibility of all proteins under the EA condition was remarkably lower than under the control condition. Using the control condition, even though digestion of soy protein in the gastric phase was faster than that of meat proteins, it was not significant different from that of meat proteins in intestinal phase. Interestingly, the digestibility of meat proteins under the EA condition, except for the soy protein, was slowly approaching the level observed for the control condition (Figure 1B). The digestibility of soy protein under the EA condition at 240 minutes (9.40%) was still far lower than that under the control condition at 120 min (35.1%). These results indicate that also in the intestinal phase digestion of soy protein was severely reduced following the gastric EA condition.

### 3.2 Dynamic digestive profiles evaluated by SDS-PAGE

The profiles of digesta during the gastrointestinal digestion process are illustrated in Figure 2. For the gastric phase, the band distribution and changes in intensity along digestion time of each meat protein group were extremely similar under both control and EA conditions. As the digestion time increased, the intensity of high molecular weight proteins diminished slightly. For example, the intensity of bands around 270 kDa and 110 kDa decreased, and that of 5 kDa increased along with time reached the maximum at 120 min. This result is consistent with Zhao et al. (2019), who investigated the digestion of beef semimembranosus proteins and found that myosin heavy chain (around 270 kDa) and α-actinin (around 110 kDa) were gradually hydrolyzed over time. It is worth mentioning that even though there were similar changes of the intensity of the protein bands under the EA condition over time, these changes appeared less pronounced as compared to digestion under the control condition (Figure 2A). For instance, the intensity of the bands around 5 kDa was reduced under the EA condition compared to the control condition (Figure 2A).

**Figure 2.**
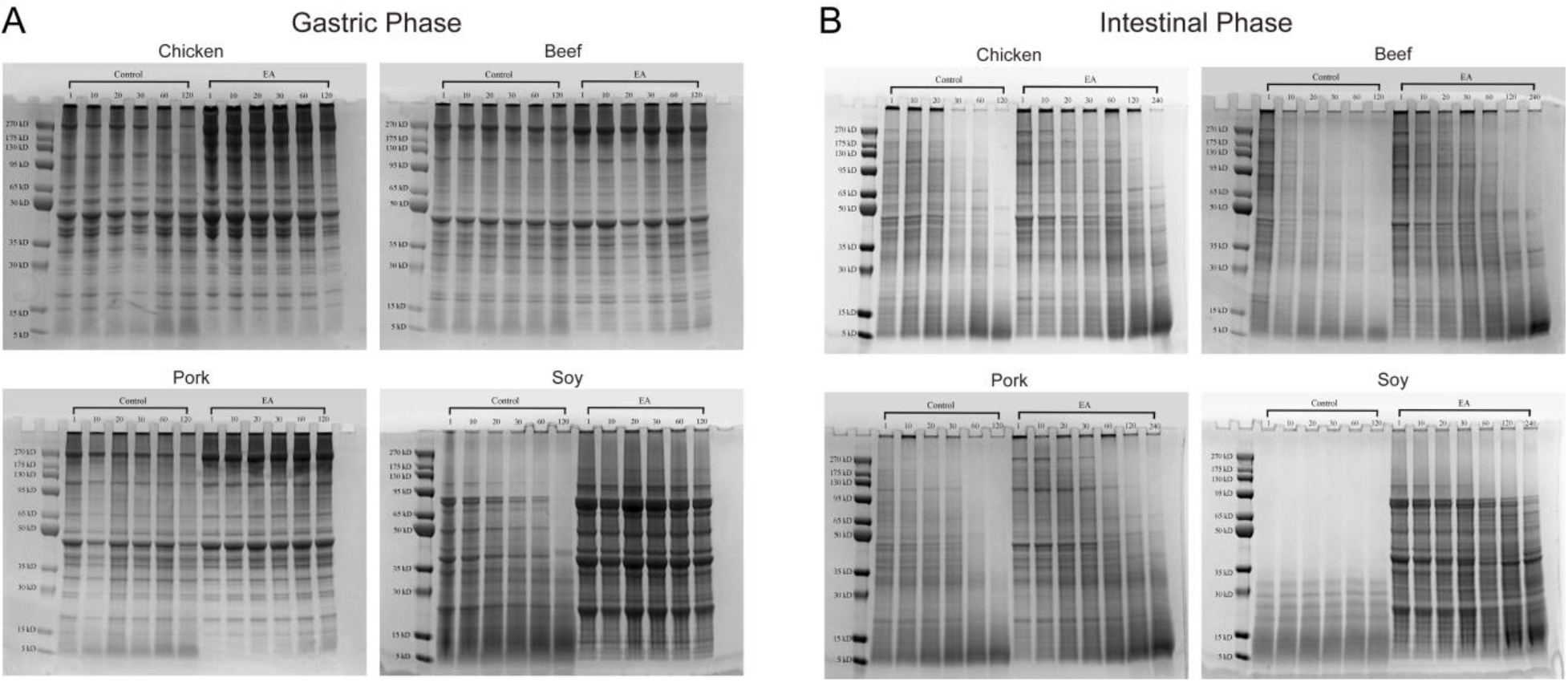
SDS-PAGE profiles of different proteins in gastric phase (A) and intestinal phase (B). Meat and soy proteins were subjected to in vitro digestion using conditions that mimicked the gastro-intestinal conditions in young healthy individuals (control, C) and elderly individuals with decreased secretion of HCl in the stomach, achlorhydria (EA). Lanes 1, 10, 20, 30, 60, 120 and 240 refer to sample digestion time (min).

Examining the SDS gel pattern of soy protein digests, we found that the difference in digestion progression between control and EA conditions was much more pronounced than observed for meat proteins. For example, the intensity of bands around 80, 50, 40, and 20 kDa decreased quickly with digestion time for soy protein under the control condition, especially for the 80 and 50 kDa bands, which almost disappeared at 120 min. By contrast, the intensities of these bands decreased very little under the EA condition. Based on previous studies, we found that these proteins comprise β-conglycinin with three subunits (α: 76 kDa, α’: 72 kDa and β: 53 kDa), glycinin with acidic polypeptide (31-45 kDa) and basic polypeptide (18-20 kDa) (Tian et al., 2019). A similar phenomenon was reported by Nguyen, Bhandari, Cichero, & Prakash (2015), showing that the intensity of the bands for β-conglycinin, acidic polypeptide and basic polypeptide decreased with increasing digestion time. In addition, in line with the studies by Yang et al. (2016), basic polypeptides of glycinin are degraded slower than acidic polypeptides which may reflect that the basic polypeptides are more hydrophobic, and thus, more compact and less accessible to pepsin.

In the intestinal phase, all proteins were further degraded both under control and EA conditions. For the meat proteins, although the digestion profiles under the EA condition was quite closed to those of the control condition, digestion still proceeded slightly slower compared to the control condition. For instance, the band around 110 kDa almost disappeared for chicken and pork under the control condition and disappeared completely for beef under the control condition at 30 or 60 min, but was clearly visible under the EA condition. Interestingly, the intensity of bands (5-15 kDa) at 120 and 240 min under the EA condition was higher than under the control condition, possibly due to continued proteolysis under the control condition yielding peptides <5 kDa migrating out of the gel. For soy protein, pronounced differences were observed between control and EA conditions in accordance with the result of DH. For soy, all high molecular weight proteins were digested into fragments with a molecular mass less than 35 kDa under the control condition at all digestion times, while they were barely digested under the EA condition. The results of digestion using the control condition were consistent with Nguyen et al. (2015), who concluded that the hydrolysis of β-conglycinin, glycinin with acidic and basic polypeptide progressed rapidly in the simulated duodenal phase and exhibited reduced intensity of the cognate bands already after 30 min of digestion. As digestion proceeded, a relatively large number of small peptides were generated with molecular masses around 20 kDa. However, of note, we observed that digestion of soy protein under the EA condition remained at a low level. Moreover, the degree of digestion in the intestinal phase of soy protein under the EA condition seemed to be even more impaired that that observed under the control condition in the gastric phase.

### 3.3 Peptide concentrations measured by Nanodrop

Due to the limitation of SDS-PAGE, peptides with a molecular mass less than 5 kDa could not be reproducibly detected on the gel. Therefore, to assess the characteristics of the digesta using different digestion conditions at the level of low molecular weight peptides, peptide concentrations were determined using Nanodrop measurements (Figure 3). Under the control condition, the peptide concentrations of all proteins increased steadily with digestion time both in the gastric and the intestinal phases. The highest peptide concentrations were observed at the final time point, corresponding to 2.87 and 3.04 mg/mL for chicken, 2.94 and 3.05 mg/mL for beef, 2.78 and 2.89 mg/mL for pork, 2.22 and 2.15 mg/mL for soy in the gastric and the intestinal phase, respectively. There was no significant difference in the peptide concentration among the three meat proteins. This was consistent with the study by Martini, Conte, & Tagliazucchi (2019), which also reported no significant differences in peptide contents obtained from beef, chicken and pork after in vitro gastro-intestinal digestion. However, the peptide concentration of soy protein was significantly lower when compared to meat proteins, which was inconsistent with the result of the DH and SDS-PAGE analyses. It has been reported that pepsin and pancreatic peptidases involved in protein digestion may catalyze the formation of short peptides and free amino acids, respectively (Heda, Toro, & Tombazzi, 2019; Wardlaw & Insel, 1996). We speculate that the lower concentration of low molecular weight peptides from digestion of soy protein could be due to the further degradation of peptides into amino acids which could not be detected by Nanodrop at 280 nm.

**Figure 3.**
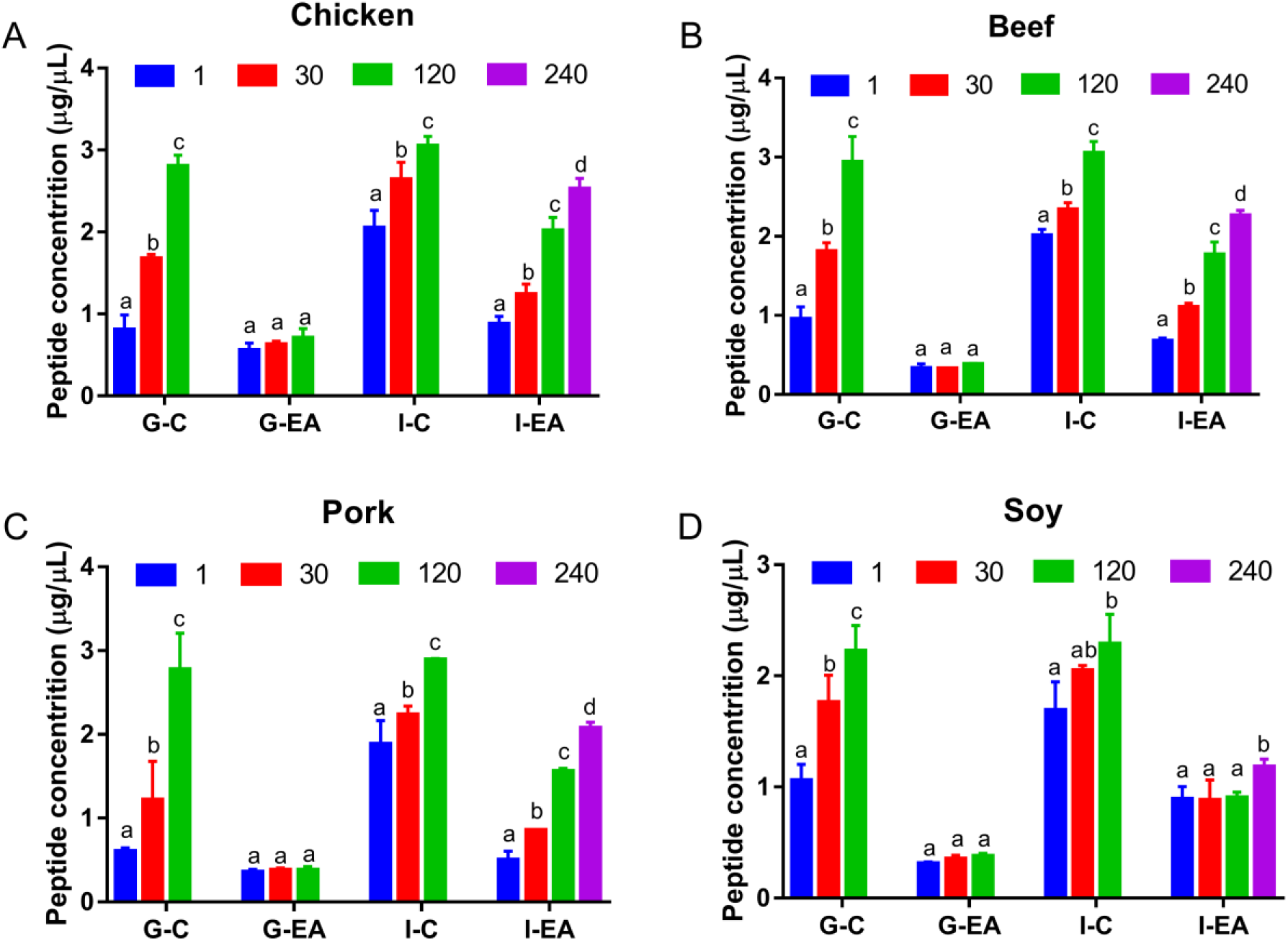
Peptide concentration of different proteins after digestion for 1, 30, 120 and 240 min. Meat and soy proteins were subjected to in vitro digestion using conditions that mimicked the gastric and intestinal conditions in young healthy individuals (G-C, I-C) and elderly individuals with decreased secretion of HCl in the stomach, achlorhydria (G-EA, I-EA), respectively. The peptide concentration was detected with Nanodrop at 280 nm. Bars with different letters are significantly different at the level *p* < 0.05. 1, 30, 120 and 240 refer to sample digestion time (min).

For the EA condition, we found that although the peptide concentration increased over time in the gastric phase, there were no significant changes. At 120 min, the peptide concentration in the chicken protein group was the highest (0.61 mg/mL), followed by beef (0.39 mg/mL), pork (0.38 mg/mL) and soy (0.32 mg/mL). This result agreed with the results of DH and SDS-PAGE. Although the DH of all proteins under the EA condition also increased over time, the overall DH was still at an extremely low level. For instance, the DH of chicken protein only increased from 0.30% at 10 min to 1.05% at 120 min. Likewise, the SDS-PAGE patterns also showed that there were almost no visible changes in the protein bands. For the intestinal phase, we found that the peptide concentration of the meat protein groups continued to increase significantly with time, with the chicken group showing the highest concentration (2.52 mg/mL), followed by beef (2.26 mg/mL) and pork (2.07 mg/mL). The peptide concentration of the meat proteins under the EA condition at 240 min was very close to that observed for control condition at 30 min. The DH results showed a similar tendency. A most noteworthy observation was that the peptide concentration of soy protein in the intestinal phase following the EA condition in the gastric phase did not show a significant difference until 240 min, which was directly related to the low degree of digestion. In addition, when we compared the overall trends of peptide concentration and DH in the meat protein groups, we observed that they were very similar. However, we observed a slight difference for the individual meat proteins. For instance, the DH of the intestinal phase of pork under the EA condition was higher than chicken and beef, but the peptide concentration of pork was lower than chicken and beef. We speculate that this may reflect the principle of the method for measuring these two indicators and the different composition among chicken, beef and pork. Overall, the digestibility of all proteins under the EA condition was lower than that under the control condition, and soy protein showed the strongest pH and enzyme concentration dependency compared to the other three meat proteins.

### 3.4 Proteomics profile during digestion

Considering the limited analytical resolution of SDS-PAGE, LC-MS/MS was applied to investigate the profiles of protein digestion products at 1, 30, 120 and 240 min. The statistics and Venn diagrams of the identified proteins are presented in Figure 4 and Suppl. Table 2, respectively. As can be seen from the table, the number of identified proteins in the chicken group was the highest (269), followed by soy (225), beef (148) and pork (80). The proteomics results of the meat protein groups were similar to the previously published study by Wen et al. (2015), who also reported that the highest number of peptides were identified in the chicken group after in vitro digestion, followed by beef and pork. Therefore, we conclude that there was a higher diversity of chicken digestion products than of beef and pork. In addition, highly abundant proteins were identified as myofibrillar proteins and sarcoplasmic proteins in the meat protein groups, including myosin, troponin C, actin and phosphorylase. These results were consistent with those reported by Li et al. (2017) and Martini et al. (2019). To examine the variations of identified protein, we calculated the percentages of shared proteins for each protein source between control and EA conditions in the gastric and intestinal phases relative to the total number of proteins (Suppl. Table 2), and found that except for the soy protein, the proportions of shared proteins in all protein groups increased from the gastric to the intestinal phase. This result revealed that the differences of meat protein digestion between EA and control conditions decreased with time, while it increased for soy protein.

**Figure 4.**
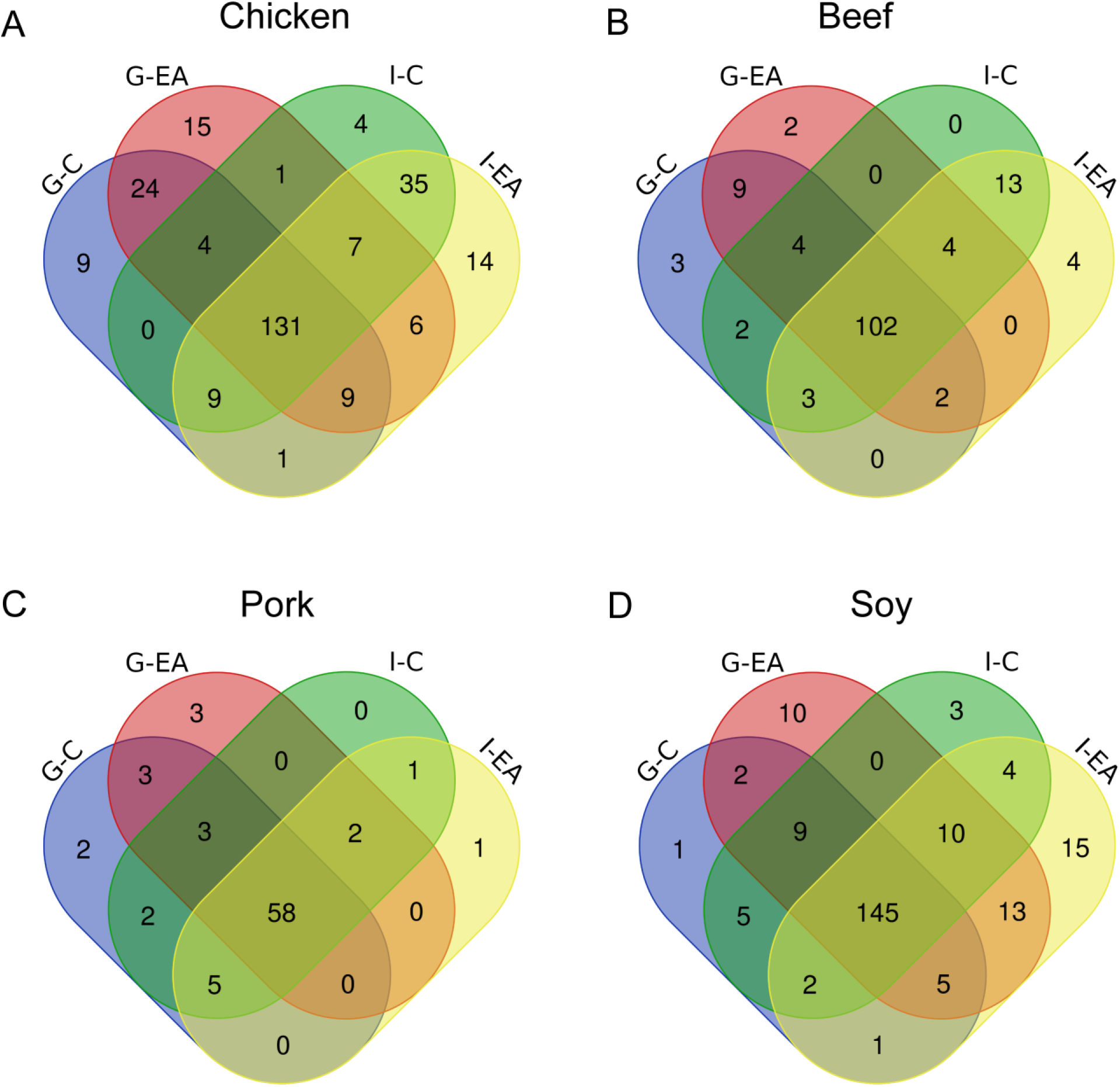
Venn diagrams of identified proteins after gastric and intestinal digestion under control and elderly with achlorhydria conditions. Meat and soy proteins were subjected to in vitro digestion using conditions that mimicked the gastric and intestinal conditions in young healthy individuals (G-C, I-C) and elderly individuals with decreased secretion of HCl in the stomach, achlorhydria (G-EA, I-EA), respectively. The value on the Venn diagrams refers to the number of identified proteins.

To evaluate the characteristics of different protein profiles at both the qualitative and quantitative levels, principal component analysis was applied based on the protein intensity (Figure 5). There was a high degree of similarity among meat protein groups. Specifically, the profiles of the gastric control (G-C) and the gastric EA (G-EA) conditions were clearly separated on both PC1 and PC2; however, the difference between intestinal control (I-C) and intestinal EA (I-EA) conditions greatly decreased. For the soy protein, G-C and G-EA, I-C and I-EA were all separated from each other, respectively. It is worth noting that the distance of soy protein between G-C and I-C conditions was very small compared to meat protein groups, which implied that soy protein was to a great extend digested in the gastric phase. These results confirmed of the results obtained using SDS-PAGE analysis, and further emphasized that the digestion of meat protein and soy protein was differentially affected by changes in the pH and pepsin concentration in the gastric phase with soy protein being more susceptible to the changes of digestion pH and enzyme concentration.

**Figure 5.**
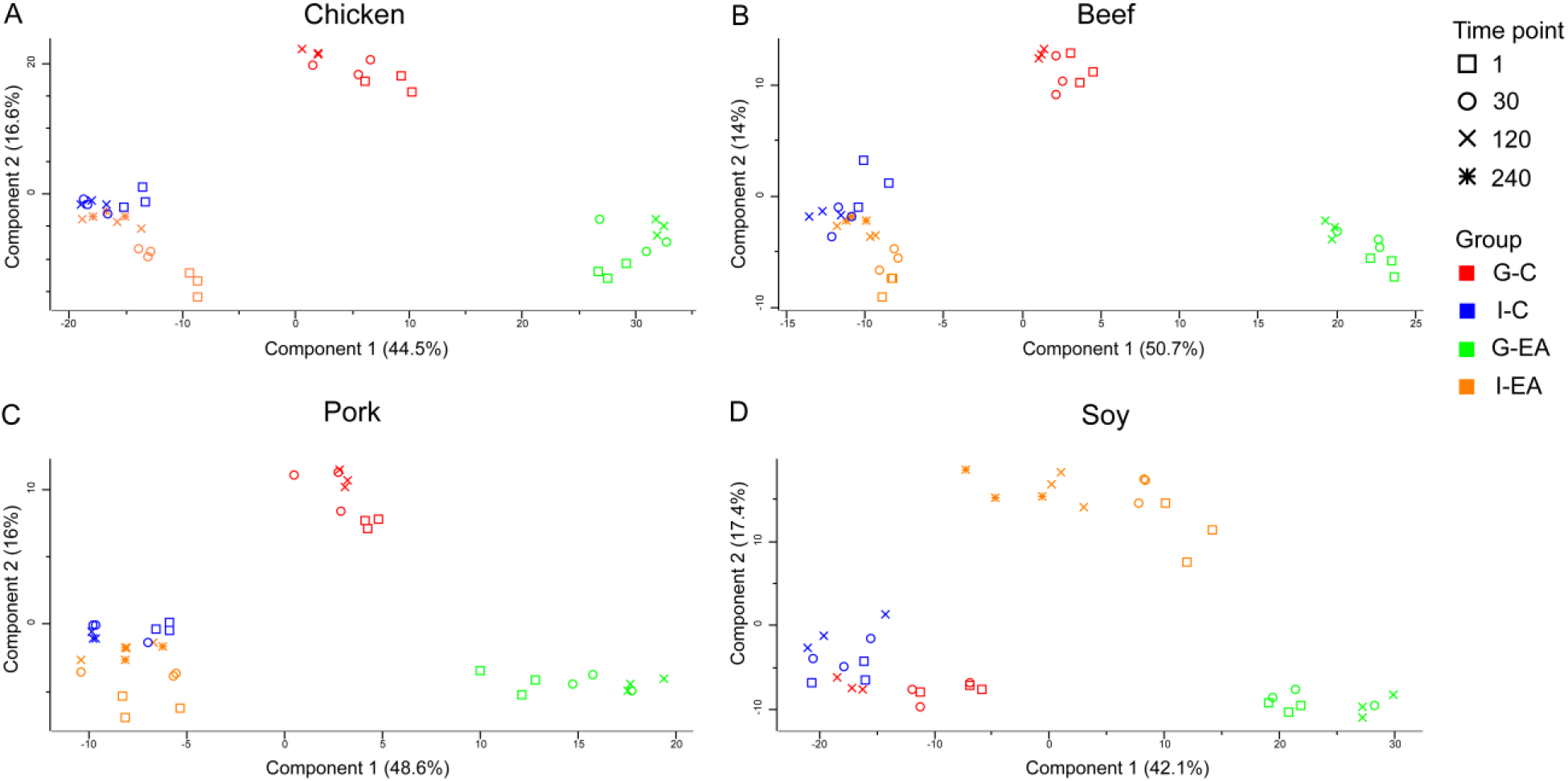
Principal component analysis for the identified proteins after gastric and intestinal digestion under control and elderly with achlorhydria conditions. Meat and soy proteins were subjected to in vitro digestion using conditions that mimicked the gastric and intestinal conditions in young healthy individuals (G-C, I-C) and elderly individuals with decreased secretion of HCl in the stomach, achlorhydria (G-EA, I-EA), respectively. 1, 30, 120 and 240 refer to sample digestion time (min).

### 3.5 Differentially abundant proteins

To further examine which specific proteins were susceptible to the changes under the different digestion conditions, we analyzed proteins that exhibited differential abundance comparing control and EA conditions. We term such protein differentially abundant proteins (DAPs) (Figure 6). We defined DAPs as proteins that exhibited an average fold change in abundance (EA/C) ≥ 1.5 or ≤ 0.67, and a *p* value < 0.05 at each time point between control and EA conditions. In total 50, 38, 24 and 67 DAPs were filtered out in the digests of chicken, beef, pork, and soy protein, respectively. For the meat proteins, the DAPs in the gastric phase accounted for a large proportion of the DAPs, a proportion that was greatly reduced in intestinal phase. However, the number of DAPs derived from soy protein did not decrease significantly going from the gastric to the intestinal phase. This result indicated that the difference in the digestion of meat protein between control and EA conditions was significant in gastric phase, but fewer differences were observed in the intestinal phase. By contrast, the difference between control and EA conditions remained high for soy protein both in the gastric (34 DAPs) and the intestinal (33 DAPs) phase.

**Figure 6.**
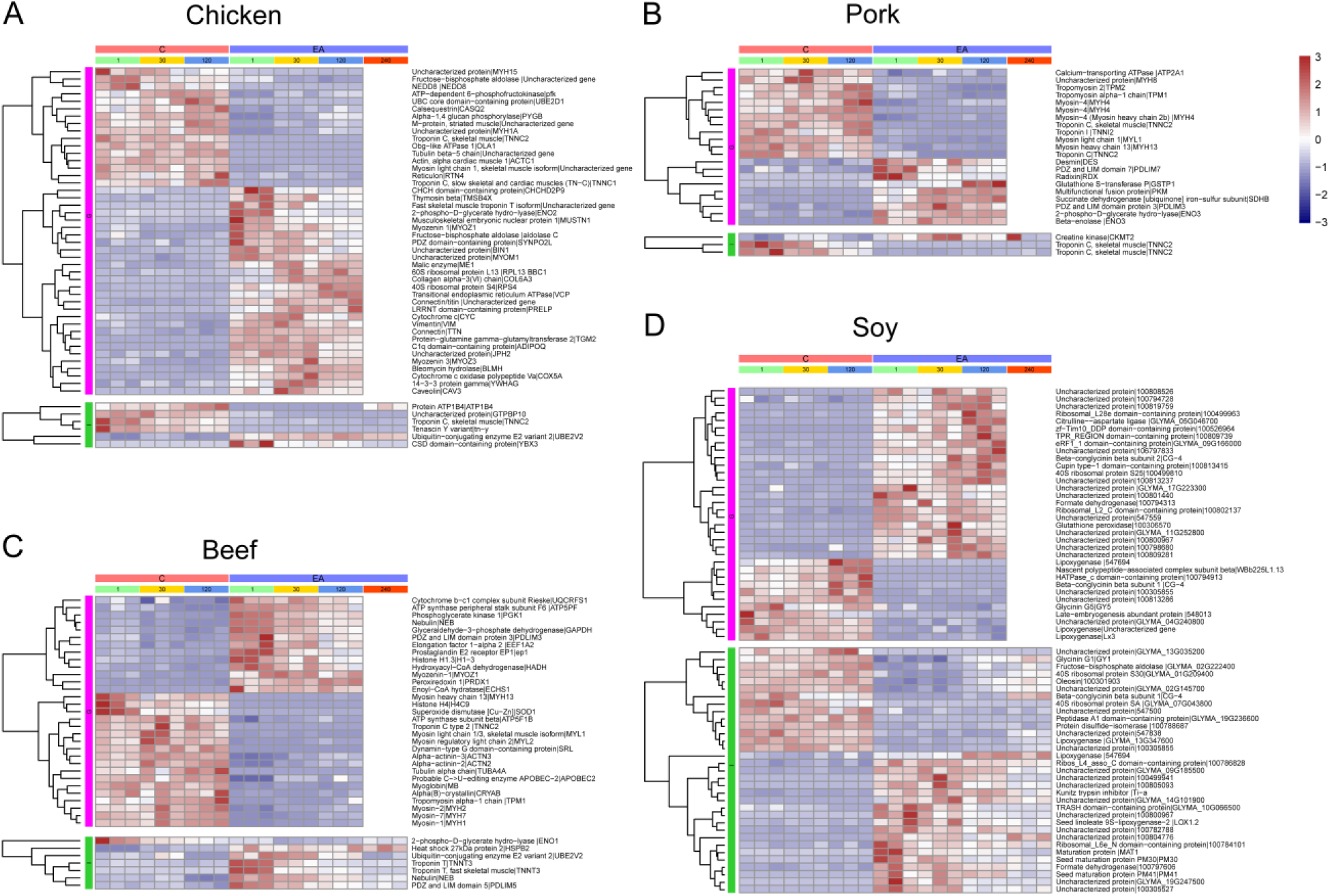
Hierarchical clustering of identified differential abundant proteins. Meat and soy proteins were subjected to in vitro digestion using conditions that mimicked the gastro (G)-intestinal (I) conditions in young healthy individuals (control, C) and elderly individuals with decreased secretion of HCl in the stomach, achlorhydria (EA). The left side of the protein description is the name of protein, the right side of the protein description is the name of the corresponding gene. 1, 30, 120 and 240 refer to sample digestion time (min).

For the meat proteins, myofibrillar proteins (myosin, actin, tropomyosin and troponin) and sarcoplasmic proteins (fructose-bisphosphate aldolase, phosphoglycerate kinase 1 and beta-enolase) were mainly identified under control and EA conditions. This result is consistent with the study showing muscle proteins as the main sources of peptides in all in vitro digested meat samples (Martini et al., 2019). In addition, the major DAPs identified with higher intensity under the control condition in the gastric phase were myofibrillar proteins, especially for digestion of pork protein, indicating a higher degree of degradation of these proteins under control conditions than EA conditions. On the contrary, the abundance of myozenin, PDZ and LIM domain protein, and sarcoplasmic proteins was higher under the EA condition. Previous studies found that myofibrillar proteins are hydrolyzed more easily than sarcoplasmic proteins during in vitro digestion (Rodríguez, Núñez, Córdoba, Bermúdez, & Asensio, 1998). The abundance of myofibrillar proteins was indeed higher than that of sarcoplasmic proteins under the control condition in our experiments (Suppl. Table 3). However, this situation was not recapitulated under the EA condition. We speculate that the increased pH might lead to a change in the structure of the protein, which led to the difference in digestibility.

For soy protein, it should be noted that due to the limitation of the UniProt database, many of the identified proteins were unannotated, while few proteins from meat were unannotated. A high proportion of unannotated proteins from soy was also reported by Koo et al. (2015), who identified 15 unknown proteins from 33 gel spots in soy. In addition, we observed that the degree of degradation of glycinin, beta-conglycinin and lipoxygenase under the control condition was higher than under the EA condition. This result is consistent with the results of the SDS-PAGE analysis. The intensity of glycinin and beta-conglycinin bands under the control condition was significantly decreased, while there were no significant changes under the EA condition. In gastric phase, taking beta-conglycinin beta subunit 1 (CG-4) as an example, the abundance was higher and increased significantly with the digestion time under the control condition (Suppl. Table 3). The change of glycinin G1 (GY1) in the intestinal phase during the digestion process was also very pronounced. GY1 was degraded at the initial time point under the control condition and the degradation gradually increased over time, while the abundance under the EA condition was comparable to control until 240 min of digestion.

In conclusion, this study systematically evaluated the effect of digestion conditions of meat proteins and soy protein in the gastric and intestinal phase employing conditions that mimicked the conditions found in healthy adults (controls) and elderly with increase stomach pH as found in individuals suffering from achlorhydria (EA conditions). Our results demonstrated that the EA GI condition markedly influenced meat and soy proteins digestion, including DH, peptide generation, and protein characteristics (SDS-PAGE, intensity and DAPs). For meat proteins, the digestion profiles were affected by the altered gastric conditions, but the differences between EA and control conditions gradually diminished in the intestinal phase. The digestion profile of soy protein was more susceptible to the altered GI digestive conditions compared to the meat proteins. These results serve as first step contributing to the establishment of dietary recommendation especially for the elderly individuals based on the possible effects of the prevalent changes in the GI tract observed in a relatively large fraction of elderly individuals.

## AUTHOR INFORMATION

### Corresponding Author

Telephone: +86 25 8439 5730.;

Telephone: +45 3532 4443.

## ACKNOWLEDGEMENTS

This research was supported by the National Natural Science Foundation of China (32001721) and Priority Academic Program Development of Jiangsu Higher Education Institutions (RAPD).

## Notes

The authors declare no competing financial interest.

