## Supplemental Table 1 for "Effect of gastrointestinal alterations mimicking elderly conditions on in vitro digestion of meat and soy proteins"

| **Supplementary Table 1. GI digestive conditions of control and elderly with achlorhydria** | | |
| --- | --- | --- |
| Digestive phase | Model | |
|  | Control (C) | Elderly with achlorhydria (EA) |
| Gastric phase | pH 3.0 | pH 6.0 |
|  | Pepsin 2000 U/mL | Pepsin 1500 U/mL |
|  | 0, 10, 20, 30, 60, 120 min | 0, 10, 20, 30, 60, 120 min |
| Intestinal phase | pH 7.0 | pH 7.0 |
|  | Bile 10 mM | Bile 5 mM |
|  | Pancreatin 100 U/mL | Pancreatin 50 U/mL |
|  | 0, 10, 20, 30, 60, 120 min | 0, 10, 20, 30, 60, 120, 240 min |
