## Supplemental Table 2 for "Effect of gastrointestinal alterations mimicking elderly conditions on in vitro digestion of meat and soy proteins"

| **Supplementary Table 2. Statistics of identified proteins from in vitro digested products of four protein sources.** | | | | | | | | | | | | |
| --- | --- | --- | --- | --- | --- | --- | --- | --- | --- | --- | --- | --- |
| Source | Total protein | Gastric phase | | | | |  | Intestinal phase | | | | |
|  |  | Total | G-C specific | G-EA specific | Shared | Percentage^1^ |  | Total | I-C specific | I-EA specific | Shared | Percentage^2^ |
| Chicken | 269 | 216 | 19 | 29 | 168 | 77.78% |  | 221 | 9 | 30 | 182 | 82.35% |
| Beef | 148 | 131 | 8 | 6 | 117 | 89.31% |  | 134 | 6 | 6 | 122 | 91.04% |
| Pork | 80 | 78 | 9 | 5 | 64 | 82.05% |  | 72 | 5 | 1 | 66 | 91.67% |
| Soy | 225 | 203 | 9 | 33 | 161 | 79.31% |  | 212 | 17 | 34 | 161 | 75.94% |
| Note: 1, Shared proteins as a percentage of total gastric phase proteins; 2, Shared proteins as a percentage of total intestinal phase proteins. | | | | | | | | | | | | |
